# Systems-level Analysis of 32 TCGA Cancers Reveals Disease-dependent tRNA Fragmentation Patterns and Very Selective Associations with Messenger RNAs and Repeat Elements

**DOI:** 10.1101/135517

**Authors:** Isidore Rigoutsos, Aristeidis G. Telonis, Phillipe Loher, Rogan Magee, Yohei Kirino, Venetia Pliatsika

## Abstract

We mined 10,274 datasets from The Cancer Genome Atlas (TCGA) for tRNA fragments (tRFs) that overlap nuclear and mitochondrial (MT) mature tRNAs. Across 32 cancer types, we identified 20,722 distinct tRFs, a third of which arise from MT tRNAs. Most of the fragments belong to the novel category of i-tRFs, i.e. they are wholly internal to the mature tRNAs. The abundances and cleavage patterns of the identified tRFs depend strongly on cancer type. Of note, in all 32 cancer types, we find that tRNA^HisGTG^ produces multiple and abundant 5´-tRFs with a uracil at the -1 position, instead of the expected post-transcriptionally-added guanosine. Strikingly, these -1U His 5´tRFs are produced in ratios that remain constant across all analyzed normal and cancer samples, a property that makes tRNA^HisGTG^ unique among all tRNAs. We also found numerous tRFs to be negatively correlated with many messenger RNAs (mRNAs) that belong primarily to four universal biological processes: *transcription*, *cell adhesion*, *chromatin organization* and *development/morphogenesis*. However, the identities of the mRNAs that belong to these processes and are negatively correlated with tRFs differ from cancer to cancer. Notably, the protein products of these mRNAs localize to specific cellular compartments, and do so in a cancer-dependent manner. Moreover, the genomic span of mRNAs that are *negatively* correlated with tRFs are enriched in multiple categories of repeat elements. Conversely, the genomic span of mRNAs that are *positively* correlated with tRFs are depleted in repeat elements. These findings suggest novel and far-reaching roles for tRFs and indicate their involvement in system-wide interconnections in the cell. All discovered tRFs from TCGA can be downloaded from https://cm.jefferson.edu/tcga-mintmap-profiles or studied interactively through the newly-designed version 2.0 of MINTbase at https://cm.jefferson.edu/MINTbase.

**NOTE: while the manuscript is under review, the content on the page https://cm.jefferson.edu/tcgamintmap-profiles is password protected and available only to Reviewers.**

**Key Points:**

- *Complexity*: tRNAs exhibit a complex fragmentation pattern into a multitude of tRFs that are conserved within the samples of a given cancer but differ across cancers.
- *Very extensive mitochondrial contributions*: the 22 tRNAs of the mitochondrion (MT) contribute 1/3^rd^ of all tRFs found across cancers, a disproportionately high number compared to the tRFs from the 610 nuclear tRNAs.
- *Uridylated (not guanylated) 5´-His tRFs*: in all human tissues analyzed, tRNA^HisGTG^ produces many abundant modified 5´-tRFs with a U at their “-1” position (-1U 5´-tRFs), instead of a G.
- *Likely central roles for tRNA*^*HisGTG*^: the relative abundances of the -1U 5´-tRFs from tRNA^HisGTG^ remain strikingly conserved across the 32 cancers, a property that makes tRNA^HisGTG^ unique among all tRNAs and isoacceptors.
- *Selective tRF-mRNA networks*: tRFs are negatively correlated with mRNAs that differ characteristically from cancer to cancer.
- *Mitochondrion-encoded tRFs are associated with nuclear proteins*: in nearly all cancers, and in a cancer-specific manner, tRFs produced by the 22 *mitochondrial* tRNAs are negatively correlated with mRNAs whose protein products localize to the *nucleus*.
- *tRFs are associated with membrane proteins*: in all cancers, and in a cancer-specific manner, nucleus-encoded and MT-encoded tRFs are negatively correlated with mRNAs whose protein products localize to the cell’s membrane.
- *tRFs are associated with secreted proteins*: in all cancers, and in a cancer-specific manner, nucleus-encoded and MT-encoded tRFs are negatively correlated with mRNAs whose protein products are secreted from the cell.
- *tRFs are associated with numerous mRNAs through repeat elements*: in all cancers, and in a cancer-specific manner, the genomic span of mRNAs that are negatively correlated with tRFs are enriched in specific categories of repeat elements.
- *intra-cancer tRF networks can depend on sex and population origin*: within a cancer, positive and negative tRF-tRF correlations can be modulated by patient attributes such as sex and population origin.
- *web-enabled exploration of an “Atlas for tRFs”*: we released a new version of MINTbase to provide users with the ability to study 26,531 tRFs compiled by mining 11,719 public datasets (TCGA and other sources).

---

Activity in recent years has been drawing increasing attention to a new group of molecules that appear to be produced at the same time as transfer RNAs (tRNAs). These molecules are referred to as tRNA fragments or tRFs and are believed to arise from both the precursor and the mature tRNAs^1-3^. For those tRFs that overlap the span of the mature tRNA, four structural categories were reported originally: 5´-tRFs, 3´tRFs, 5´-halves (5´-tRHs), and 3´-halves (3´-tRHs). In a recent analysis of hundreds of human tissues we reported a fifth structural category, the *internal* tRFs or i-tRFs that comprises numerous members expressed in high abundance^4^. In the same analysis, we also demonstrated that the identity and abundance of tRFs depends on previously unrecognized variables such as a person’s sex, population origin, and race as well as on tissue, tissue state, and disease subtype^4^. Despite these dependencies, samples from the same tissue obtained from individuals with the same sex, race and disease subtype were found to express the same tRFs and with the same relative abundances, which indicates that these molecules are constitutive^4^. More recent work showed that tRNA “halves” can be produced under stress conditions^5,6^ as well as constitutively^7-9^ and to exist in variants that are not visible by standard RNA-seq^7^.

In terms of function, tRFs have been shown to associate with Argonaute^10^ in a cell-type specific manner^4^. This indicates that at least a subset of tRFs enter the RNA interference (RNAi) pathway. In addition, tRFs have been shown to be produced differentially in response to infections^11,12^, in cancer tissues compared to normal^4,13,14^, to be affected by diet^15^, by trauma^16^, to be involved in trans-generational inheritance^17^, and to regulate translation^18^.

In summary, there is very strong evidence that tRFs: 1) represent a novel category of regulatory molecules in their own right; 2) are important in homeostasis and in disease; and, 3) warrant in-depth studies^19,20^. In this presentation, we extend our earlier work^4^ to the entirety of the TCGA collection. Specifically, we processed 11,198 cancer samples representing 32 cancer types with an emphasis on identifying *intra*and *inter*-cancer features involving tRFs. The 32 cancer types included: ACC (Adrenocortical carcinoma), BLCA (Bladder Urothelial Carcinoma), BRCA (Breast invasive carcinoma), CESC (Cervical squamous cell carcinoma and endocervical adenocarcinoma), CHOL (Cholangiocarcinoma), COAD (Colon adenocarcinoma), DLBC (Lymphoid Neoplasm Diffuse Large B-cell Lymphoma), ESCA (Esophageal carcinoma), HNSC (Head and Neck squamous cell carcinoma), KICH (Kidney Chromophobe), KIRC (Kidney renal clear cell carcinoma), KIRP (Kidney renal papillary cell carcinoma), LAML (Acute Myeloid Leukemia), LGG (Brain Lower Grade Glioma), LIHC (Liver hepatocellular carcinoma), LUAD (Lung adenocarcinoma), LUSC (Lung squamous cell carcinoma), MESO (Mesothelioma), OV (Ovarian serous cystadenocarcinoma), PAAD (Pancreatic adenocarcinoma), PCPG (Pheochromocytoma and Paraganglioma), PRAD (Prostate adenocarcinoma), READ (Rectum adenocarcinoma), SARC (Sarcoma), SKCM (Skin Cutaneous Melanoma), STAD (Stomach adenocarcinoma), TGCT (Testicular Germ Cell Tumors), THCA (Thyroid carcinoma), THYM (Thymoma), UCEC (Uterine Corpus Endometrial Carcinoma), UCS (Uterine Carcinosarcoma), and UVM (Uveal Melanoma). Lastly, where relevant, we use the NIH/TCGA designations to refer to race groups (see Methods).

## RESULTS

We discovered tRFs from all 11,198 datasets of TCGA, which we make available at https://cm.jefferson.edu/tcga-mintmap-profiles. For our analyses, we used the 10,274 datasets that were not tagged with special annotations by the TCGA consortia (see Methods). Our analyses focus only on tRFs whose sequences fully overlap a mature tRNA. These tRFs can belong to one of five structural categories (5´t-RFs, i-tRFs, 3´-tRFs, 5´-tRHs and 3´-tRHs). They can also belong to two categories (exclusive and ambiguous) based on their potential genomic origin. In terms of length, all generated tRFs range from 16 to 30 nucleotides (nt). See Methods.

### A multitude of tRFs across the 32 TCGA cancer types

We used our recently developed Threshold-seq algorithm^21^ to automatically determine a support threshold for each of the analyzed datasets. Threshold-seq adapts to a dataset’s depth of sequencing while being immune to the potential presence of outliers, making it ideal for this purpose. We report tRFs that exceeded Threshold-seq’s recommended threshold in at least one of the analyzed datasets. For the range 16-30 nt, we find a total of 20,722 distinct tRFs that exceed threshold. These tRFs comprise 1,717 5´-tRFs, 16,133 i-tRFs, 2,840 3´-tRFs, and 32 5´-tRHs. We note that fragments with lengths larger than 27 nt could be truncated versions of tRFs longer than 30 nt (see Methods). 18,453 of the 20,722 tRFs have lengths between 16 and 27 nt inclusive (= 1,395 5´-tRFs, 14,478 i-tRFs, 2,574 3´-tRFs, and six 5´-tRHs). We note that i-tRFs are abundant and very diverse, in agreement with our earlier findings^4,8,22^. Of the 20,722 tRFs, 13,904 (67%) are exclusive to tRNA space whereas the remaining 6,818 have ambiguous genomic origin. For more detailed information, see Supp. Table S1.

We also adopted the approach of the TCGA working groups and carried out NMF clustering of the datasets in each of the 32 cancer types using tRF profiles instead of miRNA profiles (see Methods). Supp. Figure S1 summarizes the results of the 288 NMF runs (SKCM samples were split into two types whereas GBM was excluded – see Methods).

### Nuclear and MT tRFs exhibit distinct and cancer-dependent profiles

In previous work, we showed differences in the length and abundance profiles of nucleus-encoded vs. MT-encoded tRFs in healthy individuals from the 1,000 Genomes Project (1KG)^4^, breast^4^ as well as prostate cancer^8^ and liver cancer patients^22^ that were not part of the TCGA initiative. These results suggest that different cancer types exhibit different distributions of nucleus-encoded and MT-encoded, respectively, tRFs. Thus, we sought to examine these profiles across all TCGA cancer types.

Figure 1A shows characteristic examples of the length distributions for nucleus- and MT-encoded tRFs and for 10 of the 32 cancer types: AAC, HNSC, LAML, OV, SKCM, THCA, TGCT, UCS, UCEC, and UVM. For a detailed distribution of the different tRF categories in each of the 32 cancer types see Supp. Figure S2 and Supp. Table S2. All distributions show abundances normalized in reads-per-million (RPM). There are evident differences in the tRFs’ structural type, lengths, nuclear vs. MT origin, and relative abundances. For example, in ACC and UVM, MT tRNAs are sources of comparatively more abundant 5´-tRFs with lengths 20, 23, and 26 nt. Analogously, 30-mer proxies from nuclear tRNAs are the most abundant species in almost all 10 cancers. Also, in SKCM and OV, nuclear tRNAs are much stronger contributors of 3´-tRFs with length 18 nt, when compared to the other eight cancer types. Figure 1B provides a global view of the structural categories (5´-tRFs, i-tRFs, and 3´-tRFs) and abundances of the populations of tRFs arising from nucleus-encoded (blue palette) and MT-encoded (red palette) tRNAs across cancer types. The Figure makes the considerable diversity of these molecules clear. Figure 1C is a Principal Component Analysis (PCA) plot based on the same data and shows clear clusters for various combinations of tRF type, length, and genome of origin, corroborating the results of Figure 1B. Note that the clusters are explained by tRF length and tRF origin. Specifically, shorter molecules (usually ≤ 23 nt) are clustered together, separately from longer ones (usually ≥ 24 nt); additionally, there is clear distinction between tRFs originating in the nucleus from those originating in the MT. These findings indicate that the nuclear and MT tRNAs produce distinctly different populations of tRFs that depend on cancer type.

**Figure 1.**
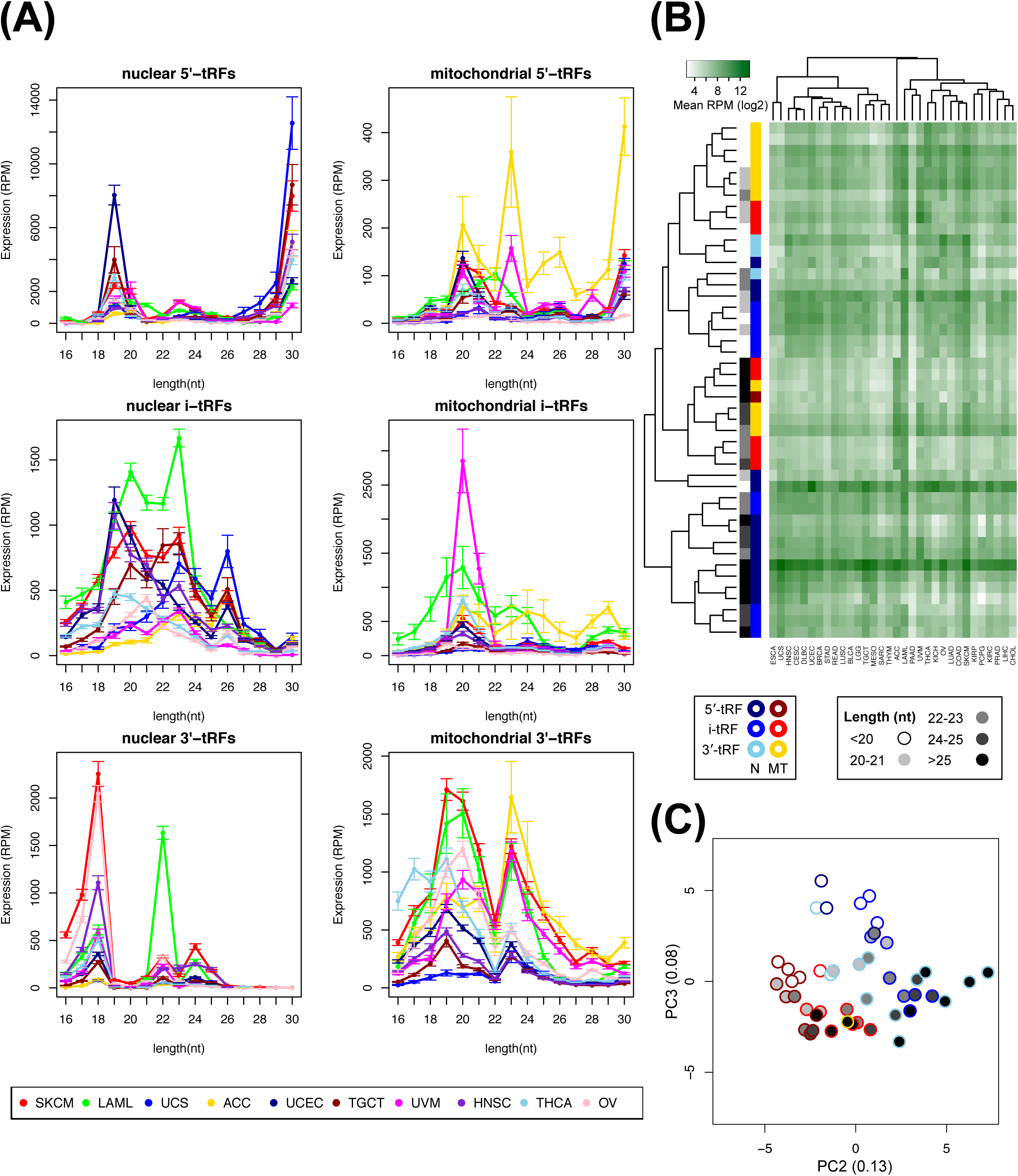
tRF distributions by category, length, and abundance. (A) Length distributions of tRFs broken down by type and organelle in 10 of the 32 analyzed cancer types (for each length the mean across samples and the standard error are shown). Each category has a unique and cancer-type-specific distribution. (B) Heatmap and hierarchical clustering (metric: Kendall’s tau coefficient) of the expression profiles of the structural categories per origin as the sum of tRF expression that fall in each category. Short tRFs (< 24nt) are clustered together in the top half of the heatmap, and separated based on their genomic origin (nucleus or MT). Longer tRFs from either nuclear or MT tRNAs are clustered together in the bottom half of the heatmap. This heatmap highlights the observation that tRFs are diverse. (C) A PCA plot showing the clustering for various combinations of tRF type, length, and genome origin, similar to the clustering shown in (B). The color-coding scheme is the same for both panels (B) and (C).

### Isoacceptors produce tRFs in a cancer-dependent manner

Having established that both nuclear and MT tRNAs are prolific producers of tRFs, we sought to investigate how different tRNA isoacceptors contribute to the abundance profiles of tRFs. We hypothesized that the production of tRFs per isoacceptor is cancer-dependent. To investigate this, we computed the expression of each isoacceptor as the sum of expression (in RPM) of the tRFs that it produces. We did so separately for each of the 32 cancer types and for each of the 61 nuclear and 20 MT anticodons. This allowed us to tag each isoacceptor with the level of expression of its tRFs on a per cancer basis. We then carried out hierarchical clustering and generated the heatmap of Figure 2A. Three isoacceptor groups are immediately evident in this Figure (indicated by the red lines). The top group comprises 21 isoacceptors producing tRFs that have lower expression on average and depend strongly on cancer type. The middle group comprises 22 isoacceptors that show moderate expression that is cancer-type-specific. The bottom group consists of 21 isoacceptors (four are from the MT) whose tRFs have high levels of normalized expression across all 32 cancer types. Three isoacceptors from this cluster, the mitochondrial tRNA^ValTAC^ and the nuclear tRNA^HisGTG^ and tRNA^GlyGCC^, stand out as producing highly abundant tRFs in all 32 cancer types.

**Figure 2.**
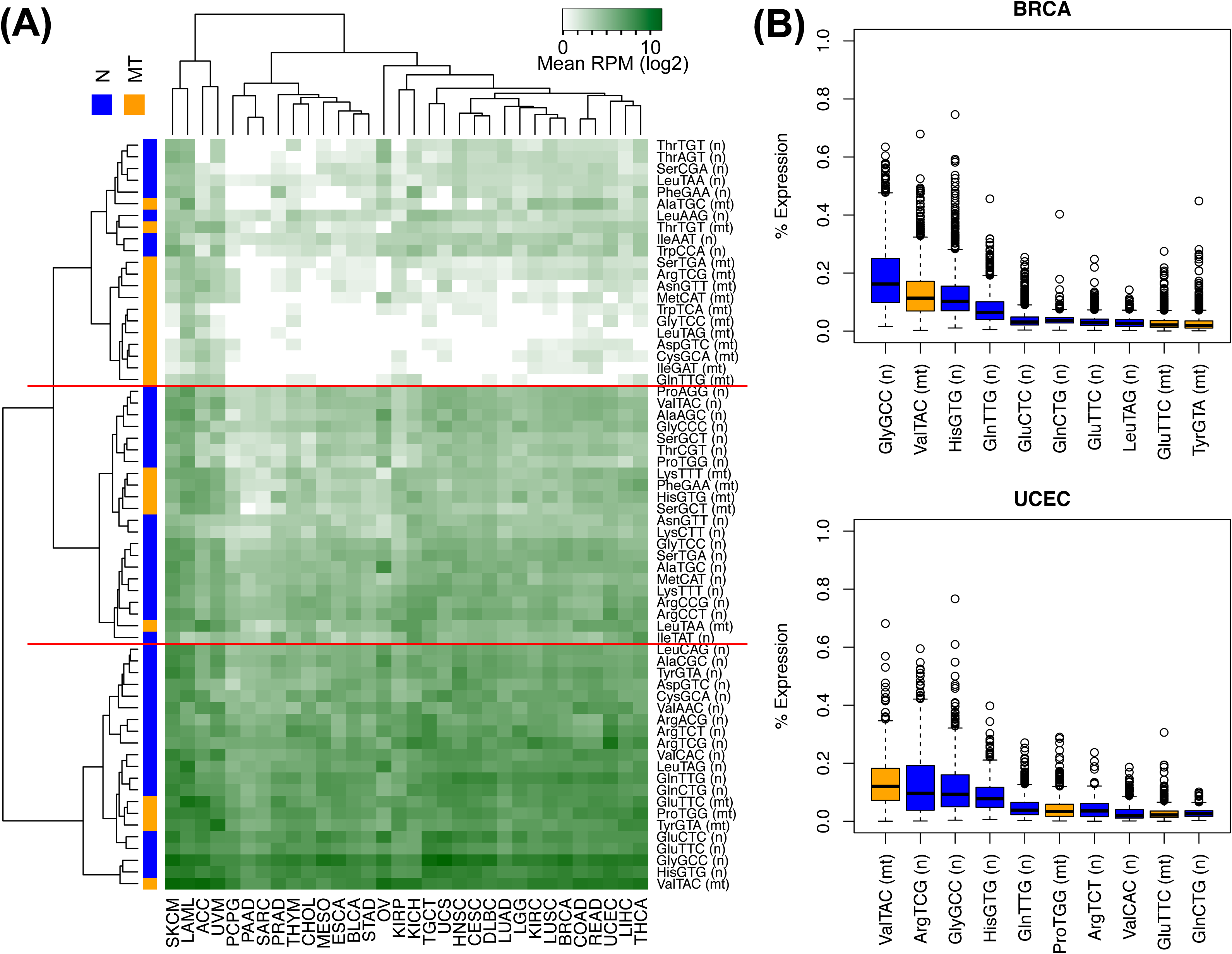
Isoacceptor representation among the tRFs. (A) Heatmap and hierarchical clustering (metric: Euclidean distance) of the abundance profile of each isoacceptor, calculated as the sum of the expression of tRFs it produces, in all 32 cancers. Nucleus-encoded isoacceptors are marked on the side color bar in blue, MT ones in orange. (B) Box-plots showing the percentage expression of tRFs from specific isoacceptors across BRCA and UCEC samples. As can be seen, the top tRF-producing isoacceptors differ in the two cancers. The highest-expressed isoacceptor in BRCA is the nuclear tRNA^GlyGCC^ whereas in UCEC it is the MT tRNA^ValTAC^ isoacceptor.

From a cancer standpoint, we note that ACC, LAML, SKCM and UVM form a distinct branch of the dendrogram (Figure 2A). All four of these cancers produce abundant tRFs from nearly all of the shown isoacceptors (see Figure 1 for comparison), albeit with significant expression variation. We highlight another example of this variation with the help of BRCA and UCEC in the boxplots of Figure 2B. In BRCA, four isoacceptors, tRNA^GlyGCC(n)^, tRNA^ValTAC(mt)^, tRNA^HisGTG(n)^, and tRNA^GlnTTG(n)^ produce most of the tRFs. On the other hand, in UCEC, it is tRFs from tRNA^ValTAC(mt)^, tRNA^ArgTCG(n)^, tRNA^GlyGCC(n)^, and tRNA^HisGTG(n)^ that are expressed abundantly. These findings indicate that the production of tRFs is cancertype-specific.

### The patterning of tRFs depends on tRF category, isoacceptor, and cancer type

In light of the results of the previous section and the dependencies of tRF abundance on cancer type and isoacceptor, we sought to identify cancer-type specific tRNA cleavage patterns. Towards this end, we analyzed “where” tRFs are located with respect to the mature tRNA’s origin. We studied all tRFs with above-threshold abundances and lengths between 16 and 27 nt inclusive. As we explain in Methods, imposing a length limit at 27 nt is unavoidable: as a result, 5´-tRHs and 3´-tRHs are not included in this analysis. To avoid contributions from tRFs of ambiguous origin, which may arise from different biogenesis processes, we focused on only the 3,136 tRFs that are exclusive to tRNA space (see Methods). In the Discussion, we discuss our findings in the context of known modifications across the span of mature tRNAs.

We tracked multiple tRF attributes and did so separately for each of the 32 cancers (see Methods). We show a holistic view of the results in Figure 3 – for the complete set of the histograms for all of the attributes see Supp. Figure S3. Note that we use a white circle to indicate the position of known modifications (m^1^G9, m^3^C32, m^1^G37, and m^1^A58) in the shown tRNA backbones^23^. We stress that we highlight these positions for reference purposes only. Indeed, it is unknown currently whether these modifications occur in the tissues and tissue states that are represented by the TCGA samples.

**Figure 3.**
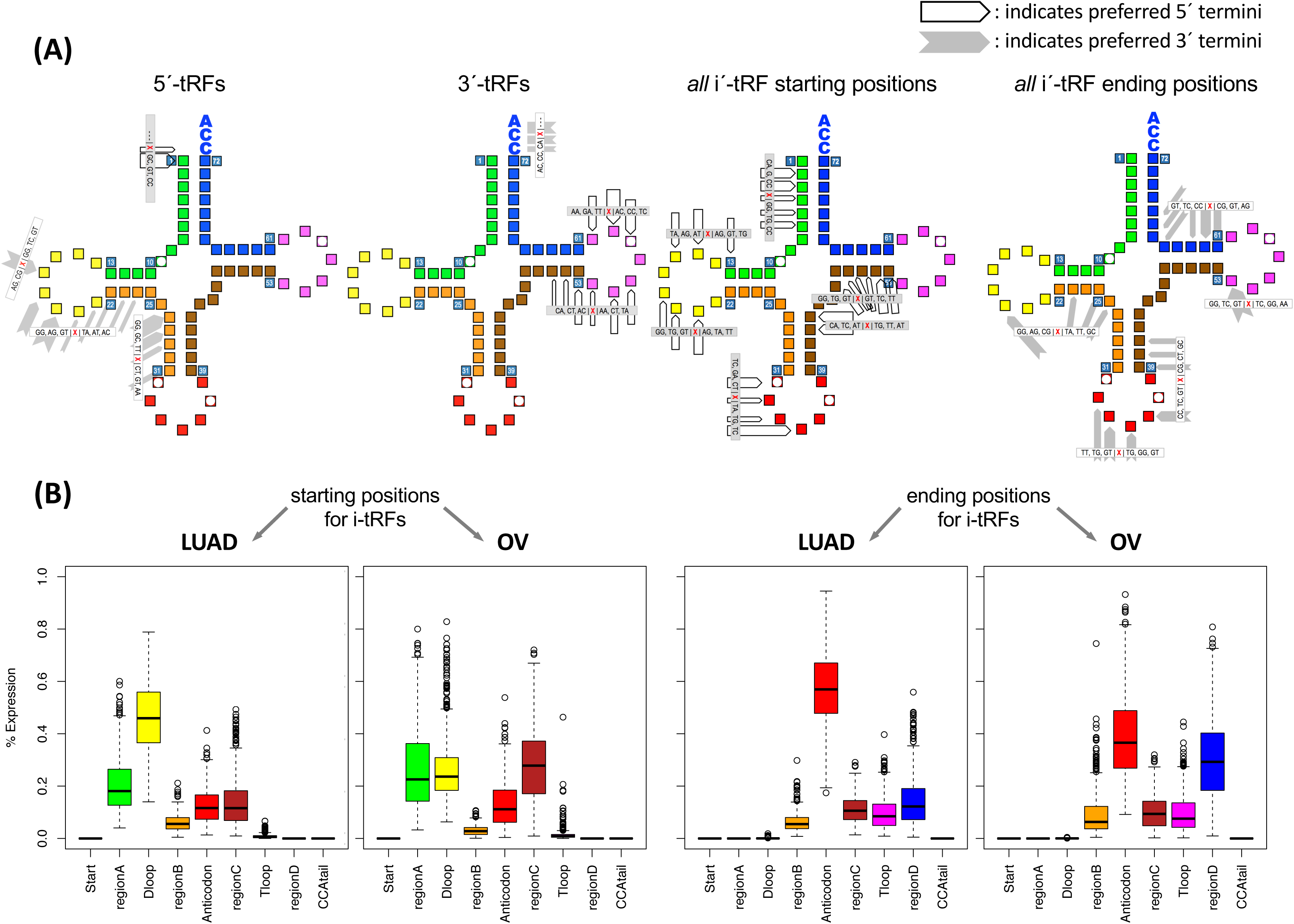
Cleavage points across the tRFs. (A) A schematic that shows the preferences of the 5´ termini (white pentagon arrows) and 3´ termini (gray chevrons) for 5´-tRFs, 3´-tRFs, and i-tRFs. For clarity purposes, separate schematics show the preferences of the 5´ termini and the 3´ termini for i´-tRFs. The thickness of the arrow or chevron indicates the preference for the corresponding position in a qualitative manner. Groups of arrows are tagged with black-on-gray labels whereas groups of chevrons are tagged with black-on-white labels. The red X of each label indicates the terminal nucleotide, either 5´ or 3´: the X is preceded (followed, respectively) by the three most frequent dinucleotides found immediately upstream (downstream, respectively) in the mature tRNA for the most abundant tRFs that begin or end at the position. Square with white circles indicate positions with known modifications. We stress that these modifications are shown for reference purposes only as it is unclear whether they occur in the tissues and tissue states that are represented by the TCGA datasets we analyzed. (B) Box plots showing the preferences for the starting (left) and ending (right) positions for i-tRFs in LUAD and OV.

In Figure 3A we see that the more abundant 5´-tRFs have only moderate preference for the location of their 3´ termini, which span virtually all positions from the middle of the D-loop through the beginning of the anticodon loop. Analogously, for the 3´-tRFs, which can terminate at any of the three nucleotides of the non-templated “CCA” addition, their 5´ termini begin just before or within the T-loop. We note that the observed preferences across human cancers for the 3´ termini of 5´-tRFs, and for the 5´ termini of 3´tRFs respectively, match the preferences that were recently reported for tRFs in the plant *A. thaliana*^24^.

Among the various fragment categories, i-tRFs are the most diverse in both their 5´ and 3´ termini choices, as we reported recently^4^. In theory, i-tRFs can begin and end at every nucleotide of the mature tRNA, except for its 5´ end or the CCA tail. However, our detailed analysis revealed that i-tRFs exhibit distinct cleavage patterns in individual cancers. As seen in Figure 3A, i-tRFs start either close to the 5´ end of the tRNA, at the D or the most 5´ half of the anticodon loop or between the variable and the T loop. The 3´ ends of the i-tRFs also favor specific positions.

We highlight the i-tRF endpoint preferences by examining in more detail the i-tRFs in LUAD and OV (Figure 3B). In LUAD, comparatively more i-tRFs begin inside the yellow region (D-loop) than do in OV. On the other hand, more i-tRFs begin in the brown region (region C) in OV than do in LUAD. Analogous comments can be made about the i-tRFs’ ending positions in LUAD and OV. See also Supp. Figure S3 for more details.

These findings indicate that the manner in which tRFs are cleaved from the respective tRNA depends on the cancer type, on the isodecoder, and on the structural type of the tRF. The findings also argue strongly against the tRFs being random products of tRNA degradation.

### Uridylated His(-1) tRFs are abundant in human tissues and exhibit a unique property that is not affected by tissue or tissue state

In eukaryotes, before the mature tRNA^HisGTG^ can be recognized by its cognate aminoacyl tRNA synthetase, guanylation of its 5´-terminus by the enzyme THG1 (THG1L in human) is required^25-27^. This post-transcriptionally added nucleotide is referred to as the “-1” position and denoted “His(-1).” In recent work with a cell line (the breast cancer model BT-474), it was shown that full-length mature tRNAs and 5´-tRHs from tRNA^HisGTG^ also contain a uracil at the His(-1) position^28^. To the best of our knowledge, this possibility has not been examined before in human tissues. We therefore sought to profile the His 5´-tRFs and the identity of their -1 nucleotide across all 32 TCGA cancer types.

Our analyses reveal that, in human tissues and across all 32 cancer types, the largest portion of 5´tRFs from tRNA^HisGTG^ contains a uracil at the His(-1) position – we will refer to them as “-1U 5´-tRFs.” A smaller fraction of 5´-tRFs contain an adenine at the His(-1) position, whereas 5´-tRFs with a guanine or cytosine are even fewer. The -1U 5´-tRFs are exclusive to tRNA space and thus can only be produced by isodecoders of tRNA^HisGTG^. However, it cannot be stated with certainty whether these -1U 5´-tRFs arise from cleavage of the precursor or from post-transcriptional modification of the mature tRNA: four of the 12 isodecoders (the one from MT and the three nuclear tRNA-His-GTG-1-6, tRNA-His-GTG-3-1, tRNAHis-GTG-1-5) contain a T at that location of the DNA template.

Even though the biogenesis of these -1U 5´-tRFs remains elusive, we found their presence in the numerous TCGA RNA-seq datasets and in a cell line^28^ intriguing and set out to study their profiles. Examination of -1U 5´-tRFs from tRNA^HisGTG^ across all 32 TCGA cancer types uncovered a striking property for those -1U 5´-tRFs that differ by a single nucleotide in their 3´ termini and have lengths between 16 and 24 nt inclusive. In particular, we discovered that as the length of these -1U 5´-tRFs increases, their abundance alternates from low to high to low to high, etc. Specifically, we discovered that the ratio of abundances of these increasingly longer fragments remains constant in all 32 TCGA cancers. Curiously, the pattern of relative abundances was the same for both the normal and the cancer state. Moreover, we found that the pattern is not exhibited by *unmodified* 5´-tRFs, i.e. by 5t´-RFs that begin at position +1 of the mature tRNA^HisGTG^, to which we refer as +1G 5´-tRFs. Figure 4 shows the log_10_ of the mean ratio of (abundance of -1U 5´-tRF ending at position i) / (abundance of -1U 5´-tRF ending at position i+1), for BLCA, ESCA, PAAD, BRCA, LUAD, and SKCM. For comparison purposes, the Figure also shows the ratio for the +1G 5´-tRFs that end at consecutive positions: as can be seen, +1G 5´-tRFs do not exhibit the pattern. If normal samples are available, we report values for both the tumor (red) and normal (green) samples. For the +1G 5´-tRFs the curves are colored gray (normal) and black (tumor). The points of the green (red, respectively) curve are shifted slightly to right (left, respectively) along the X-axis in order to make the details of both curves visible simultaneously. Similarly, the gray and black curves are shifted as well.

**Figure 4.**
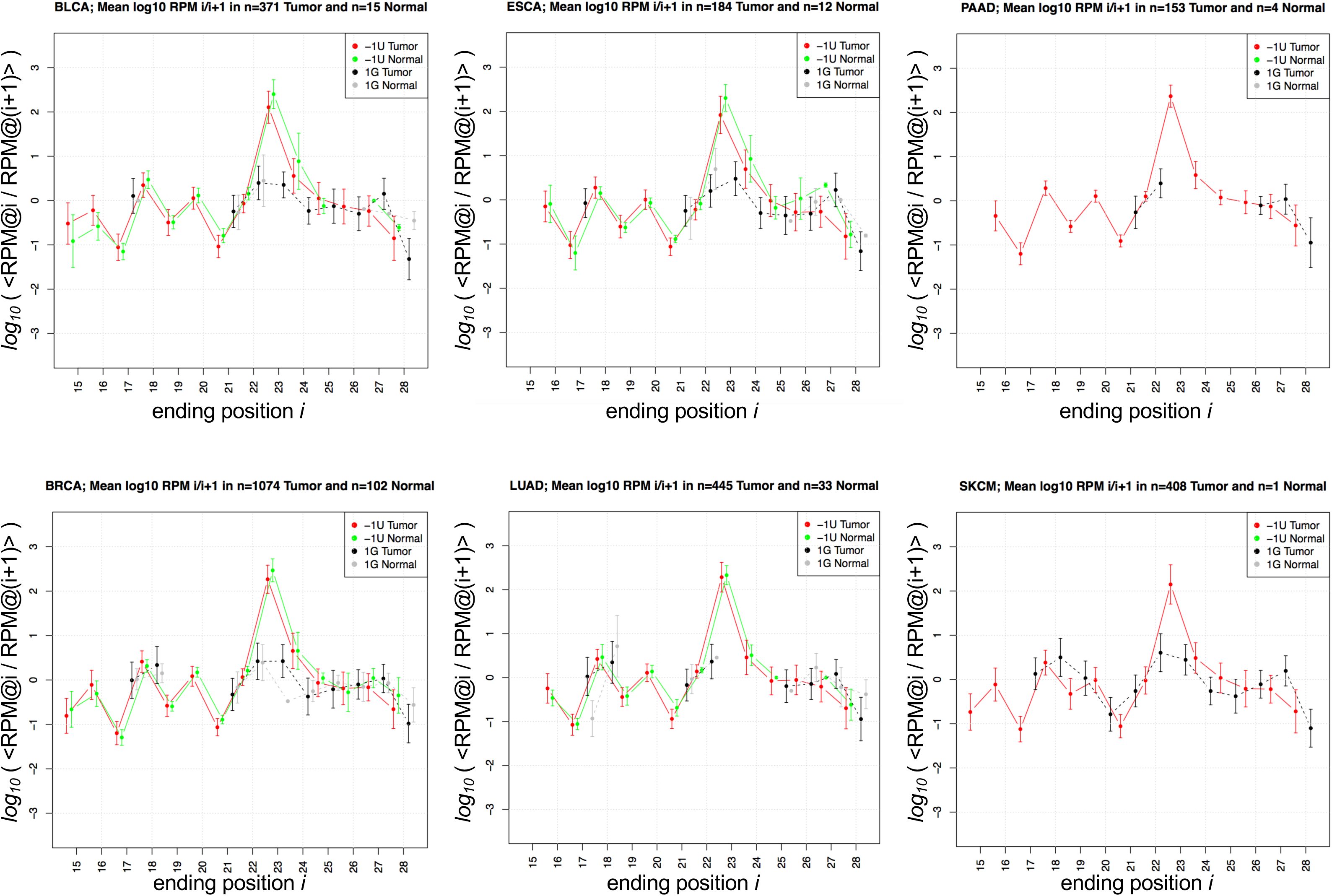
His(-1U) fragments. Abundance ratios of uridylated His(-1) 5´-tRFs from nuclear tRNA^HisGTG^ that end at consecutive positions within the mature tRNA. The shown ratios for normal (green) and cancer (red) samples represent 2,635 tumor and samples from six TCGA cancers: BLCA, ESCA, PAAD, BRCA, LUAD, and SKCM. Values are shown only for statistically significant tRFs. Y-axis: log_10_. At X=I, we plot the ratio “log_10_ (mean [(RPM of 5´-tRF ending at position *i*) / (RPM of 5´-tRF ending at position *i*+1)]).”

This finding suggests that the biogenesis of uridylated His(-1) 5´-tRFs is under exquisite control and that the specifics of this process are conserved in both health and disease, and across tissues. This conserved relationship suggests that these -1U 5´-tRFs, whether instigators or effectors, participate in cellular process that are common to all cancer types, and, thus, of essential nature. The complete collection of these plots for all 32 cancers can be found in Supp. Figure S4.

### System-level Networks: tRFs are positively- and negatively-correlated with one another in a selective manner

As part of the above analyses, we compiled the profiles of tRFs for all 32 cancer types. In our previous work, we found that tRFs from the same anticodon can be clustered in groups that are explained by the position with respect to the mature tRNA and by their lengths (see Figure 3 of Telonis *et al*^4^). Here, we expand the analysis to systematically study the correlation patterns among tRFs. For each cancer type, we computed pair-wise correlations (Spearman) between tRFs. We only kept tRF-tRF pairs whose correlation value was ≥ 0.333 or ≤ -0.333 and the associated false discovery rate (FDR) was ≤ 0.01. Multiple tRFs satisfied these criteria in each of the 32 cancer types.

Analysis of the resulting correlations revealed that the correlated tRFs exhibit notable properties that pertain to the organelle in which the tRFs are produced, the source isoacceptor, the length of the tRF, and the structural type of the tRF (Supp. Table S3, see Methods for details on how the probability values in the table were calculated). Specifically, we found the following:

– the expressed tRFs remains essentially the same across the 32 analyzed cancer types (Supp. Figure S5A);
– the expressed tRFs that participate in tRF-tRF pairs are characteristically cancer-specific (Supp. Figure S5B);
– tRFs that are positively correlated with one another originate almost exclusively in the *same* cellular compartment (either both pair members are nuclear tRFs, or, both are MT tRFs);
– tRFs that are negatively correlated with one another originate in different compartments (i.e., one of the tRFs comes from the nucleus and the other from the MT);
– positively-correlated *nuclear* tRFs frequently arise from *distinct* isoacceptors;
– positively-correlated *MT* tRFs frequently arise from the *same* isoacceptor;
– negatively-correlated tRFs frequently arise from *distinct* isoacceptors, irrespective of whether they originate in the nucleus or the MT;
– positively-correlated tRFs frequently have *similar* lengths (length difference < 5 nt) and belong to the *same* structural category;
– negatively-correlated tRFs frequently have *different* lengths (length difference ≥ 5 nt) and belong to *different* structural categories.

The complete list of tRF-tRF pairs (both positively- and negatively-correlated) and their corresponding correlation values and statistical significance can be found in can be found in the Supp. Table S4.

These results indicate that tRFs are a considerably heterogeneous group of molecules. tRF characteristics, such as isoacceptor of origin, organelle of origin, length and structural type are important determinants of the types of correlations in which they participate. We stress that, despite these commonalities, the choice of which expressed tRFs participate in positively- or negatively-correlated pairs depends on cancer-type.

### System-level Networks: tRFs are positively- and negatively-correlated with mRNAs and pathways in a selective manner

From a functional standpoint, others^10^ and we^4^ have shown that tRFs can be loaded on Argonaute, just like miRNAs. In fact, such loading was demonstrated to affect the abundance levels of mRNAs^29^. To compare and contrast the potential impact of miRNAs and tRFs in the cancer context, we leveraged the available *long* RNA-seq data of the TCGA repository. As a positive control case, we included miRNAs in these analyses. Specifically, we computed all correlated tRF-mRNA and miRNA-mRNA pairs, and examined their properties across and within cancer types. These analyses were carried out with the understanding that, for both miRNAs and tRFs, these anti-correlations capture both *direct interactions* and *indirect relationships*. The complete list of tRF-mRNA pairs (both positively- and negatively-correlated) and their corresponding correlation values and statistical significance can be found in the Supp. Table S4.

First, we examined whether tRF-mRNA and miRNA-mRNA anti-correlations persist across cancer types. As we computed abundance correlations, we were strict when filtering tRFs to minimize the inclusion of noise in our data. We found that the expressed tRFs, miRNAs, and mRNAs are essentially the same across cancers (Supp. Figures S5A, S6A, and S6B). However, what changes dramatically from one cancer type to the next is the specific manner in which miRNAs and tRFs “partner” with mRNAs to form negatively-correlated pairs. This point is evidenced by the very low off-diagonal support in Supp. Figures S5B, S6C and S6D. Within a cancer, tRFs and miRNAs are frequently negatively correlated with the same mRNAs, as evidenced by the 2x2 mini-matrices across the diagonal in Supp. Figure S6E. This suggests possible synergistic activities by miRNAs and tRFs.

Next, we examined whether the cancer-specificity of the negatively-correlated tRF-mRNA and miRNA-mRNA pairs translate into differences in the underlying pathways. To investigate this possibility, we performed DAVID analysis for each collection of mRNAs in tRF-mRNA pairs in search of enriched Gene Ontology (GO) terms and also KEGG pathways. We observed that the distribution of GO Biological Process (BP) terms as well as of KEGG pathways resembles a power-law distribution with many pathways found uniquely in one cancer-type and relatively fewer pathways appearing in several types (Supp. Figure S7A-B). For example, “renal cell carcinoma” was found enriched among the mRNAs that are negatively correlated with tRFs in KIRC. However, other enriched KEGG pathways, such as “pathways in cancer” and “proteoglycans in cancer,” are universal. Overall, we observed that any two cancers have a smaller overlap in terms of enriched mRNAs (Supp. Figure 7C) compared to enriched pathways (Supp. Figure S7D). This suggests that, although the tRF-mRNA or miRNA-mRNA correlations are cancer-type-specific, the processes that are negatively correlated with tRFs and miRNAs are more general.

We then focused on the GO terms for Biological Processes (BP) that we found to be common to multiple cancer types (Supp. Figures S7A and S7B), grouped them into non-redundant clusters (Supp. Figure S7D), and identified four main pathways: (a) Transcription, (b) Development and morphogenesis (abbreviated as “Development”), (c) Chromatin organization, and, (d) Cell adhesion and extracellular matrix organization (abbreviated as “Cell adhesion”). Notably, mRNAs from the “Transcription” and “Chromatin organization” pathways were negatively correlated predominantly with tRFs and exhibited these correlations across the vast majority of cancer types. On the other hand, mRNAs from the other two pathways (“Development” and “Cell adhesion”) were negatively correlated with miRNAs, with tRFs or both (Supp. Figure S7D).

Having established the conserved relationship between these four pathways and the associated tRFs, we examined how often tRFs overlapping isodecoders of a specific isoacceptor are associated with mRNAs from the respective GO term. Fig. 5A shows in heatmap form the fraction of cancer types in which tRFs overlapping a shown isoacceptor are negatively correlated with mRNAs belonging to each pathway.

We see that most tRNA isoacceptors are linked with the same GO terms in many cancer types. The frequency of those correlations does not depend on the tRNA’s genome of origin (mitochondrial vs. nuclear) or the encoded amino acid. We also observe that tRFs from several mitochondrial and nuclear isoacceptors are very often negatively correlated with almost all examined GO terms. The mitochondrial ValTAC, LeuTAA and ProTGG isoacceptors are negatively correlated with mRNAs from all shown GO categories (Fig. 5A) in nearly all cancer types: this is true even for pathways whose mRNAs do not have a previously reported mitochondrial link, e.g. “cell adhesion.”

Collectively, the above results provide further support to the view that the tRF-mRNA anticorrelations are an integral component of the molecular physiology of cancer, and not random. In fact, the analysis shows that tRF-mRNA anti-correlations parallel miRNA-mRNA anti-correlations (Fig. 5A). It is important to note that the tRF-mRNA anti-correlations comprise tRFs from both the nucleus and the mitochondrion, which in turn indicates that the nuclear and MT genomes marshal the corresponding pathways in a cooperative manner.

### System-level Networks: tRFs are linked to the cellular destinations of proteins encoded by negatively-correlated mRNAs

Spurred by the numerous statistically significant links between tRFs and miRNAs and pathways, we sought to examine one more facet of these associations, namely the cellular localization of the protein products of the corresponding mRNAs. For this analysis, we treated nuclear tRFs separately from mitochondrial tRFs. We used information from the UniProt database to distinguish among the following six “compartments:” nucleus, cytoplasm, endoplasmic reticulum or Golgi, mitochondrion, cell membrane, secreted, and “other organelle” (e.g., vesicles and endosomes). To evaluate the non-randomness of the localization distributions of the encoded proteins, we performed Monte-Carlo simulations to investigate the possibility of enrichments or depletions in the observed values as compared to values expected by chance.

First, we examined tRFs. The left-most and middle panels of Fig. 5B show the sub-cellular localization and distribution of the protein products of mRNAs that are negatively correlated with nuclear and MT tRFs, respectively. Several observations can be made readily. Perhaps most prominent is the finding that for several cancer types, many mitochondrial tRFs are negatively correlated with mRNAs whose protein products localize primarily to the nucleus, the cytoplasm, or the cell membrane (more than 50% in almost all cancers). On the other hand, the nuclear tRFs are negatively correlated with many mRNAs whose protein products localize to the mitochondrion (adjusted p-val < 10^-3^). COAD and SARC represent two extreme cases in this analysis. In COAD, anti-correlations involving nuclear tRFs are essentially absent. In SARC, we observed the opposite: almost no anti-correlations involved mitochondrial tRFs. One additional observation is that although in absolute numbers many proteins localize to the nucleus, cytoplasm and cell membrane, there is considerable cancer specificity as to whether this localization differs from chance, and whether the difference corresponds to *enrichment* or *depletion*. For example, in cancer types KIRC, MESO, UVM, ESCA and BLCA the nuclear tRFs are correlated with the same number of mRNAs that produce the nuclearly-localized proteins. However, in MESO and BLCA, this number is significantly lower than expected, in KIRC the number is higher, and, in UVM it is not significant. There are also cancer types in which the nuclear and mitochondrial tRFs have markedly different behavior: in SKCM and PAAD, the nuclear tRFs are negatively correlated with mRNAs coding for cell membrane proteins. For both cancer types, this number is significantly lower than expected (purple). On the other hand, in SKCM and PAAD, mitochondrial tRFs exhibit the opposite trend: they are negatively correlated with significantly more mRNAs that code for cell membrane proteins than is expected by chance. These data further highlights the cancer-specific nature of tRF profiles, their associated roles and their diversity.

We repeated the same analysis for miRNAs and show the results in the right-most panel of Fig. 5B. Similarly to tRFs, the miRNAs are negatively correlated with mRNAs whose protein products are destined for <u>all cell compartments</u> but, notably, and similarly to tRFs, more than 50% of these proteins are localized in the nucleus, the cytoplasm, and the cell membrane. However, miRNAs do not show the pronounced dependence on cancer type of tRFs (left and middle panels of Fig. 5B) as in most cases the cell membrane and secreted proteins are enriched, while nuclear proteins are usually, but not always, depleted in the respective gene sets.

These results provide strong support to the view that the observed tRF-mRNA pairs are not accidental. In fact, they resemble the results we obtain when we analyze miRNA-mRNA pairs. Thus, it is reasonable to posit a possible cooperation between miRNA and tRFs, with the miRNAs capturing the ‘base-layer’ and the tRFs overlaying a ‘cancer-dependent’ component on it. Equally importantly, the findings suggest strong associations between nuclear and mitochondrial tRFs with proteins that operate beyond the nucleus and the MT compartments.

### System-level Networks: the genomic span of mRNAs that are positively- or negatively-correlated with tRFs are selectively enriched/depleted in specific repeat elements

In light of our earlier work^30-32^ and the more recent findings in mouse that connect fragments from tRNA^GlyGCC^ with the MERVL repeat and mRNAs^15^, we hypothesized that a link between tRFs and repeat elements exists in human cancers.

We focused on all of the mRNAs that we found to be statistically significantly correlated with tRFs. We analyzed these mRNAs separately for each cancer type. For each cancer type, and for each of RepeatMasker’s^33^ categories of repeat elements, we determined the fraction of these tRF-correlated mRNAs that corresponded to fragments from the repeat category at hand. In each case, we evaluated whether the observed fraction of embedded fragments was expected by chance. We achieved this by running 10,000 iterations of a Monte-Carlo simulation that allowed us to assign a z-score to the fraction (see Methods). Compared to chance, positive z-scores represented enrichment in this repeat category’s sequence fragments. Analogously, negative z-scores represented depletion. We analyzed sense instances of repeat element fragments separately from antisense ones. Also, we analyzed positively-correlated mRNAs separately from negatively-correlated ones.

Figure 5C shows a heatmap of the generated z-scores, for all 32 cancer types, for sense and antisense instances of all repeat categories, and, separately for mRNAs that are positively correlated or negatively correlated with tRFs. The very high or very low z-scores strongly argue that these findings are not random. As Figure 5C makes apparent, in all 32 cancer types, the genomic spans of mRNAs that are either positively or negatively correlated with tRFs exhibit significant enrichment or depletion in repeat elements or their reverse complements.

**Figure 5.**
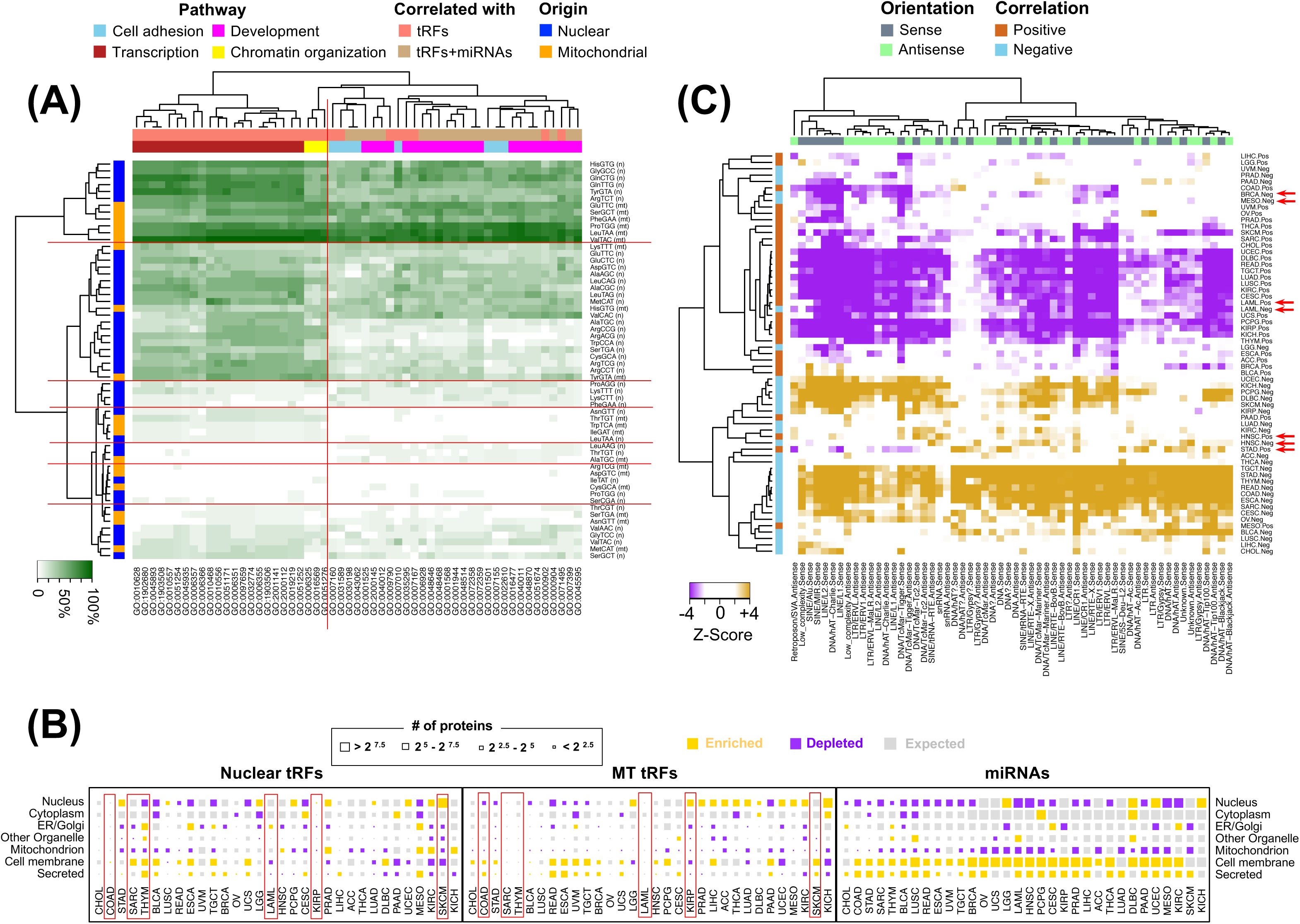
Correlations, Compartments and Repeats. (A) Heatmap and hierarchical clustering (metric: Euclidean distance) depicting the fraction of the 32 cancer types in which the shown 58 isoacceptors (rows) are anti-correlated with the listed GO terms (columns). The descriptions for the shown terms appear in Supp. Table S6. The same isoacceptors correlated negatively with the same pathways, but not at the gene or tRF level across cancers (Supp. Figure S3). Thin red lines have been added to facilitate the elucidation of the various groupings. (B) The localization of the protein products whose mRNAs are statistically significantly anti-correlated with tRFs, and mRNAs. The size of the block corresponds to the number of protein products that localize in the shown compartment. The color of the block represents enrichment (yellow) or depletion (purple) compared to the expected distribution (p-val < 0.01). A gray colored block indicates no deviation from the expected distribution. Red rectangles highlight cancers showing distinct differences in the nuclear and MT heatmaps. (C) Heatmap and hierarchical clustering (metric: Pearson correlation) showing the statistical significance (z-score) of the enrichment or depletion of fragments from repeat categories in the genomic loci of mRNAs that are anti-correlated with tRFs.

From an mRNA standpoint, we observe that the genomic spans of mRNAs that are *negatively correlated* with tRFs are *enriched* in multiple categories of repeat elements, in both sense and antisense orientation. Conversely, we observe that the genomic spans of mRNAs that are *positively correlated* with tRFs are *depleted* in repeat elements. For example, in READ, SKCM, KIRP, TGCT, and PCPG, those mRNAs that are positively correlated with tRFs are *depleted* in L1, L2, and ALU elements. In the same five cancer types, those mRNAs that are negatively correlated with tRFs are *enriched* in L1, L2, and ALU elements. A handful of the 32 cancers are notable exceptions to this observation: STAD, HNSC, LAML, BRCA, and MESO (indicated by arrows in Figure 5C).

Considering that many tRFs have repeated genomic instances, it is possible that the correlations we observe are the result of ambiguous tRFs whose multiple genomic instances outside of tRNA space overlap with mRNAs. We examined all possible genomic origins of such tRFs and could not find support for this hypothesis (Methods and Supp. Figure S9).

These results provide additional independent support to our earlier findings that the distribution of repeating sequences in the human genome is not arbitrary^30-32,34^. Moreover, the uncovered associations between tRFs and repeat elements strongly implicate the latter in the layer of tRF-mediated regulation of expression in nearly all 32 cancer types.

### System-level Networks: Intra-cancer networks of tRFs can be modulated by a patient’s sex or a patient’s race

We hypothesized that tRF profiles differ across sex or race boundaries and investigated the matter in two cancer types for which sex-dependent and race-dependent disparities of genetic origin, respectively, have been documented in the literature. Spurring this hypothesis is the above finding that tRFs are strongly associated with tRFs, mRNAs, and proteins that localize to specific cellular compartments.

Before proceeding further, we mention that in the below analysis we limit ourselves to only two of the 32 cancers types contained in TCGA. Additional in-depth studies that escape the scope of this presentation will be necessary in order to examine whether the analysis of RNA-seq datasets from other TCGA cancer types supports similar findings. We stress here that mining RNA-seq data is distinctly <u>*unlike*</u> the task of detecting, e.g., race-based somatic *mutations*, for which TCGA is well known to be underpowered^35^.

The first of the two cancer types is LUAD. In lung cancer, both sex and race disparities are known to exist. A portion of these disparities can be attributed to differences in the stage and degree of adoption of tobacco smoking^36-41^. However, age-adjusted lung cancer incidence rate is higher among black men compared to white. Also, it is roughly equal between black and white women, even though black men and black women have a lower overall exposure to cigarette smoke. These observations suggest that sex and race contribute to these differences^42^. Below, we examine only the sex-dependence aspect of LUAD.

The second cancer type is the subtype of BRCA known as “triple negative” (TNBC). TNBC represents approximately 15-20% of the BRCA cases^43^ and is the most aggressive BRCA subtype, characterized by poor prognosis. In the absence of an expressed hormone receptor, chemotherapy continues to remain the only systemic option for TNBC patients^44^. TNBC is twice as frequent among B/Aa premenopausal women compared to Wh women^44-49^.

In each case, we formed networks of tRFs whose expression values were statistically significantly correlated: we only kept relationships with a Spearman correlation ≥ 0.33 or ≤ -0.33 and a matching false discovery rate (FDR) ≤ 0.05. Then, we examined whether and how these networks changed between males and females in LUAD and between White and Black/African American patients with TNBC.

#### Case: LUAD

We analyzed the lung adenocarcinoma samples from TCGA separately for male and for female patients. The top row of Figure 6 show the network of *negatively* correlated tRF pairs that satisfy the correlation value and FDR thresholds mentioned above and are supported by the LUAD samples in TCGA. The next two rows show the subset of edges and vertices that correspond to tRF-tRF correlations that are exclusive to *male* (2^nd^ row) or *female* (3^rd^ row) LUAD patients. The 4^th^ row shows those tRF-tRF correlations that are present in both *male* and *female* LUAD patients. The networks are colored based on which mature isoacceptor produces the tRFs (1^st^ column), the tRFs’ structural category (2^nd^ column), the tRFs’ lengths (3^rd^ column), and whether the tRF originates in a mature tRNA from the nucleus or the MT (4^th^ column). As can be seen, female LUAD patients exhibit more and more-widespread anticorrelations compared to male patients.

**Figure 6.**
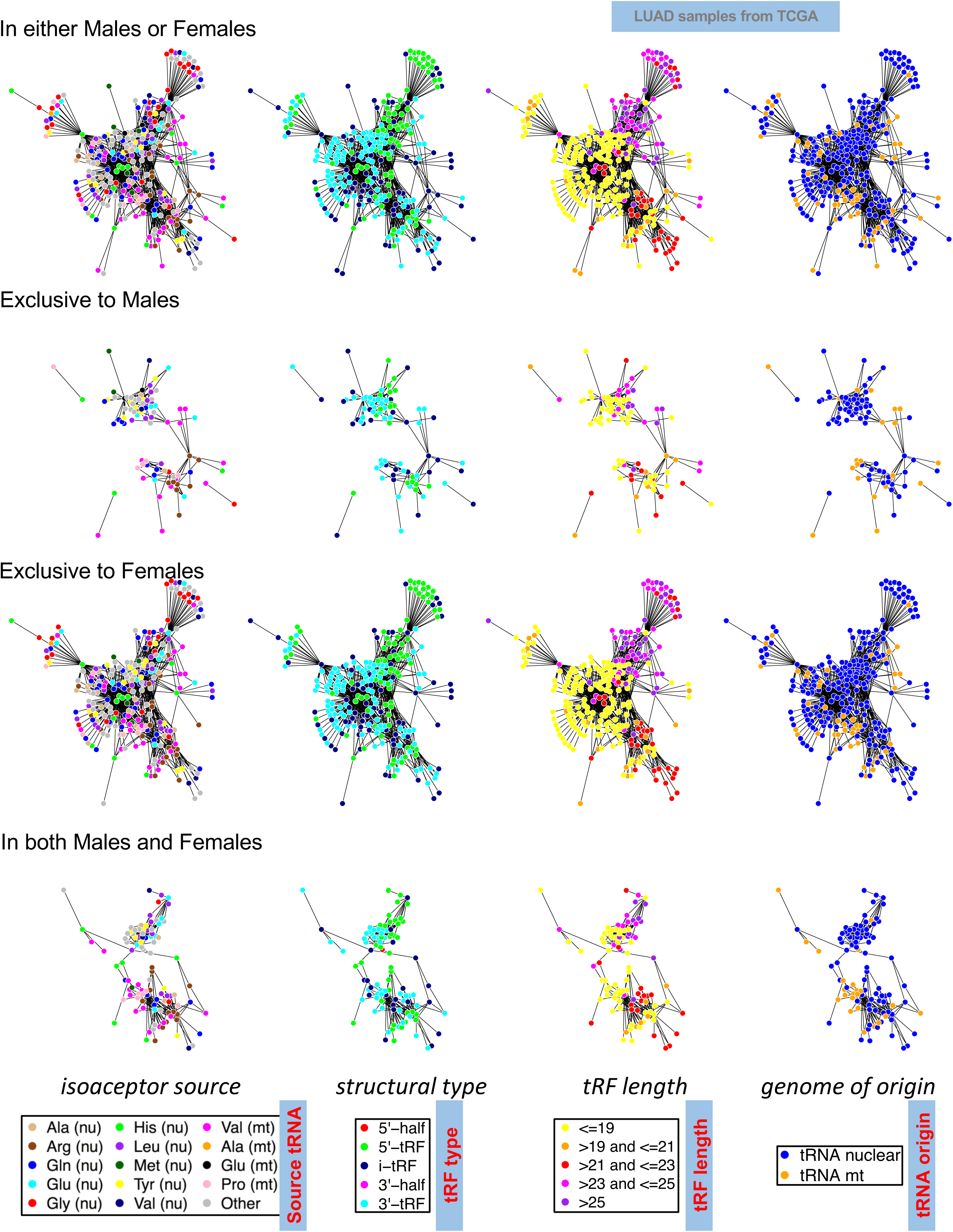
tRF correlations in patients of different sex. Example showing the dependence of the tRF profiles on the sex of patients with lung adenocarcinoma. Shown are the networks of tRF-tRF correlations that are supported by all LUAD samples from TCGA, the sub-network of correlations that are present exclusively in samples from male LUAD patients, the sub-network of correlations that are present exclusively in samples from female LUAD patients, and, finally, the sub-network of correlations that are present in LUAD patients of both sexes. From left to right, the networks are color-coded by source tRNA, structural category, length, and nuclear/MT origin. Edges between nodes correspond to a Spearman correlation ≤ -0.5 (negative correlations) and have an associated FDR ≤ 0.01. See also text.

#### Case: TNBC

We analyzed the TNBC samples from TCGA and created analogous networks. Here, it is the networks of *positively*-correlated tRF pairs that show characteristic differences between White (Wh) and Black/African American (B/Aa) patients with TNBC (see “Nomenclature/Notation” in Methods). Supp. Figure S8 shows the network of tRF-tRF pairs for all TNBC patients, the subset of the network that is present only in Wh TNBC patients, only in B/Aa TNBC patients, and, in both Wh and B/Aa patients. As in the case of LUAD, there are evident differences in the networks of correlations that are present in the Wh and B/Aa TNBC patients, respectively.

### The discovered TCGA tRFs can be studied using a newly-added MINTbase module

We recently reported the development of MINTbase, a framework for storing and studying tRNA fragments^22^. MINTbase is both a web-based content repository and a tool for the interactive study of tRFs. Originally, we populated MINTbase with 7,129 unique and statistically significant tRFs that resulted from our analyses of 832 public datasets^4,8,9,22^.

We have now extended MINTbase (version 2.0) to include the tRFs that we generated in our analyses of TCGA. With the addition of the tRFs from 32 TCGA cancer types, MINTbase now comprises information about the location, normalized abundances, and expression patterns of 26,531 distinct tRFs compiled by mining a total of 11,719 public datasets from TCGA and elsewhere.

To extend the utility of the repository, we augmented its search capabilities. Specifically, we now allow the user to search using a TCGA cancer abbreviation (e.g. BRCA, PRAD, PAAD, etc.), a descriptive phrase (e.g. breast cancer), one or more structural categories, one or more isoacceptors, a sequence (e.g. GGCTCCGTGGCGCAATGGA), a tRNA name, or a tRNA label, and to combine these choices with a “minimum abundance” criterion. As an example, the following complex Boolean request can be executed by pointing-and-clicking: “retrieve <u>all 5´-tRFs</u> and <u>all i-tRFs</u> that overlap with either <u>the *mitochondrial* isodecoder of tRNA^AspGTC^</u> or <u>any of the *nuclear* isodecoders of</u> tRNA^HisGTG^ and are <u>present in any of the breast cancer samples</u> of MINTbase with <u>abundance ≥ 25 RPM</u>.”

Each of MINTbase’s 26,531 tRFs has its own exclusive record that lists all publicly known identifiers for it, information about the isodecoder(s) that contain it, a multiple sequence alignment in the case of multiple tRNA origins, whether the tRF is exclusive to tRNA space^4,8,9^, and how many of the MINTbase datasets contain the tRF with an abundance of ≥ 1.0 RPM.

To enable *intra*-TCGA comparisons as well as comparisons between TCGA and non-TCGA datasets, each tRF record includes four histograms that show: the *fraction* of datasets containing the tRF in each TCGA cancer type and outside TCGA; the tRF’s *distribution of abundances* in each TCGA cancer type and in non-TCGA datasets; and, two more histograms showing box-plots of the distribution of abundances of the tRF *within* each TCGA dataset using a linear and a log_2_ Y-axis, respectively. All four histograms are interactive and allow the user to select which dataset(s) to display. In Figure 7, we show three of the four histograms from the record of the -1U 5´-tRF from tRNA^HisGTG^ with sequence TGCCGTGATCGTATAGTGGTT. The top histogram shows that the tRF is present in at least 75% of the samples that are available for 31 of the 32 TCGA cancer types. The only exception is LAML where the fragment appears in only 29 of the 191 datasets. Of the 521 non-TCGA datasets currently contained in MINTbase, the fragment is present in only 8 of them. Across the TCGA datasets in which it is present, this -1U 5´-tRF exhibits a wide range of abundances that reach as high as 1,394.78 RPM in LIHC (not shown). To demonstrate the comparative differences of the fragment’s distribution of abundances, we selected and show the histogram bars for COAD, LUSC, PAAD, PRAD, SKCM, UCEC and UVM (middle panel). For the same set of cancer types, in the Figure’s bottom panel, we also show the box-plot of their abundance distributions (note that this panel uses a log_2_ Y-axis). To facilitate inclusion in user reports, all these diagrams can be saved in PNG, JPG, PDF or SVG format (Figure 7, middle panel).

**Figure 7.**
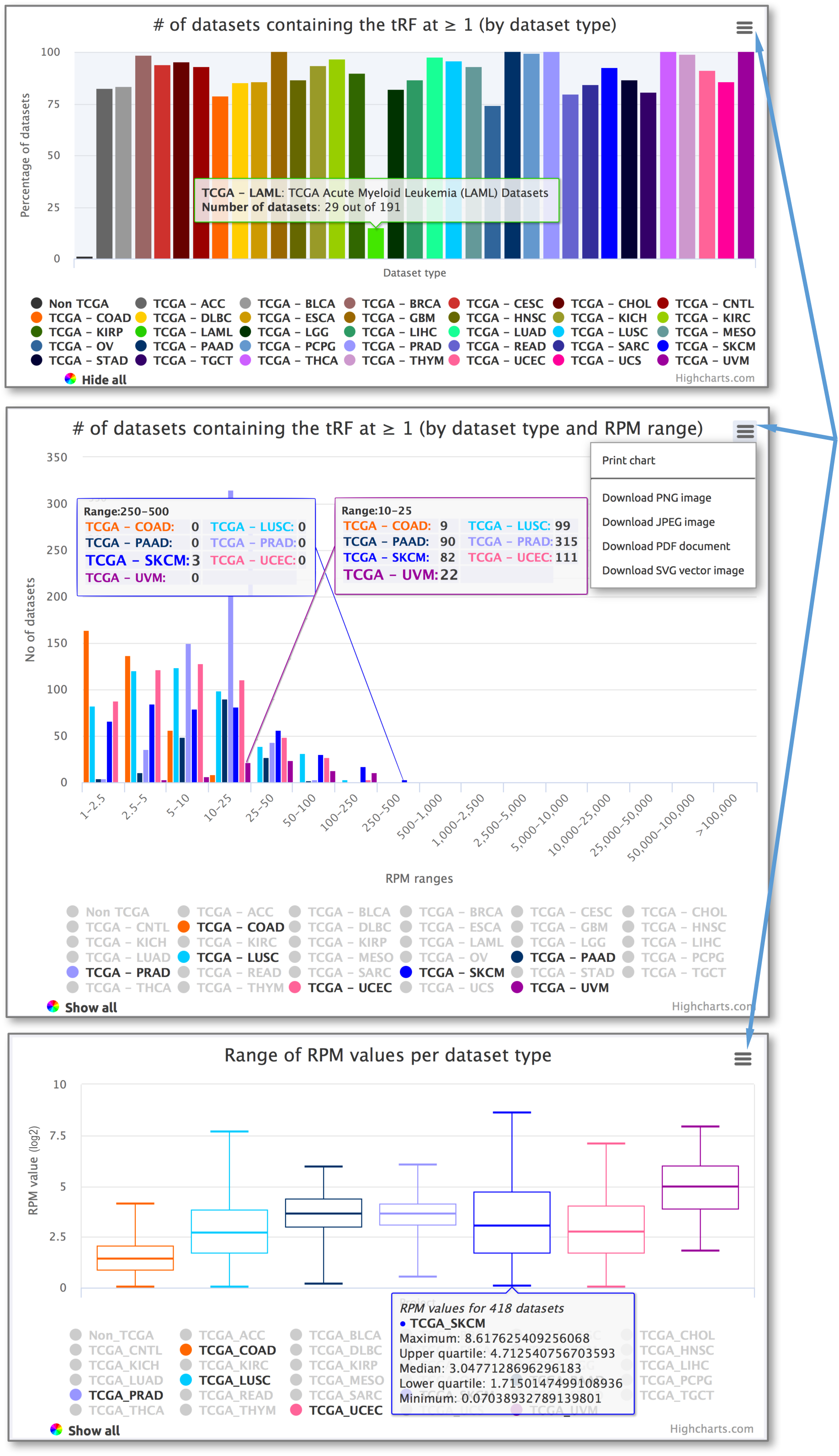
v2.0 of MINTbase. We have augmented the interface of MINTbase to enable interactive and detailed exploration of the tRFs contained in it. Here we show three of the four histograms and other information that is available in the record of the His(-1) 5´-tRF TGCCGTGATCGTATAGTGGTT from tRNA^HisGTG^ (note the starting “T”). See text for more details.

## Discussion

We carried out a comprehensive mining of 11,198 datasets from TCGA in search of statistically significant tRNA fragments. 10,274 of these datasets representing 32 human cancer types had associated records that are devoid of any special annotations (“whitelisted”) and entered our downstream analyses. We found that nearly all tRNAs exhibit cancer-specific cleavage patterns. Additionally, we found that nucleus-encoded and MT-encoded tRNAs exhibit distinctly different behavior vis-à-vis to patterning and the abundance of the tRFs they generate. tRNA^HisGTG^ represents an exception in that it gives rise to a specific collection of 5´-tRFs that contain a uracil in their -1 position (instead of the expected guanosine). The relative abundances of these -1U 5´-tRFs exhibit ratios that are maintained constant across all examined cancer types and in both health and disease. The analyses also revealed wide-ranging associations between tRFs on one hand, and mRNAs and proteins on the other. Many of the (positive and negative) associations involve partners that cross organelle boundaries: for example, they involve tRFs that arise from nucleus-encoded tRNAs and mRNAs whose proteins localize in the MT; or, tRFs that arise from MT-encoded tRNAs and mRNAs whose proteins localize in the nucleus. These associations provide new insights to understanding the layer of post-transcriptional regulation. Moreover, in the short term, these relationships suggest intriguing novel viewpoints from which to study inter-organelle communication. In the longer term, there is great potential in leveraging these relationships to develop novel diagnostics and novel therapeutics that are tuned to individual cancers.

We note that we carried out our study with full understanding that the presence of documented modifications across the span of tRNAs^50-56^ has the potential to pause or stop reverse transcription. Such modifications would result in some, perhaps many, tRFs to not be represented among the sequenced reads^57,58^. Two recently reported methods^57,58^ introduced a demethylation step that was shown to improve the enumeration of tRFs for certain isoacceptors. It is thus highly probable that the 20,722 tRFs we have identified are but a subset of a larger class of tRFs that are active in cancer tissues. By studying the TCGA datasets, we work with the best and most comprehensive datasets that are available for the time being. Even though these datasets are arguably incomplete, they remain invaluable in helping us shed a first light on important characteristics of tRFs across tissues.

A key finding of the analysis is the diversity of the identified tRFs. We mined a total of 20,722 tRFs that range widely in terms of abundance, length, structural type, and the location of their 5´ and 3´ termini. The tRFs also depend on the identity of the corresponding template isodecoder/isoacceptor^10,59^ and the identity of the genome (nuclear vs. mitochondrial) hosting the tRNAs from which the tRFs arise^22^. The type of the cancer that is analyzed each time further modulates these tRF attributes; we will return to this point below. Given that our computational pipeline complements other available methods by exhibiting superior sensitivity and specificity^60^, our results significantly enrich the publicly available data with new information that can be readily exploited in future studies.

Approximately one third of all identified tRFs are of ambiguous origin. In other words, if one examines the entirety of the human genome, the sequences of these tRFs can be found within annotated tRNAs as well as at loci that contain only *partial* instances (e.g., one half or one third) of mature tRNA sequences^4,9,22^. Some of these loci resemble full-length tRNAs^4,61^ whereas other loci correspond to partial tRNAs, repeat elements or mRNAs^4^, and, possibly, non-transcribing sequences. Recognizing this complication, in parallel work, we designed and implemented MINTmap^60^, a freely-available tool that facilitates the identification of tRFs of ambiguous genomic provenance. Strictly speaking, ambiguous tRFs require special attention, particularly when experimental work is being considered, as they cannot be linked unequivocally to transcription from a tRNA template. We provided examples of non-exclusive tRFs that are correlated with mRNAs containing an embedded instance of the corresponding tRF. Even though there were few such examples in TCGA, they warrant caution because their biogenesis may not be linked to tRNA transcripts.

The 22 mitochondrial tRNAs were found to be very strong contributors to the pool of distinct tRFs, when compared to the 610 nucleus-encoded tRNAs. In fact, 30% of all discovered tRFs derive from the 22 MT tRNAs (Supp. Table S1). This finding mirrors our previous results^4,8,9,22^ and extends them to the numerous human tissues that are part of TCGA. Moreover, MT-tRNA-derived tRFs show marked differences when compared with the nuclear-tRNA-derived tRFs. Indeed, for a given cancer type, the mitochondrial tRFs differ from their nuclear counterparts in length, relative abundances, dominant structural category, etc.

Even when we confine ourselves to a specific genome, i.e., nuclear or MT, we find a strong dependence of the tRF populations on the identity of the parental isoacceptor. These populations change across cancer types and are characterized by differences in the structural type of the produced tRFs (Figure 1), the identity of the isoacceptor that produces most distinct tRFs (Figure 2), and the relative abundances of the tRFs (Supp. Table S3). Of the 32 cancers, SKCM, UVM, ACC and LAML rank highest in their richness in distinct and abundant tRF populations.

Moreover, the tRF populations show cancer-dependent differences with regard to the specific endpoints that are favored by tRFs of a given structural type (Supp. Table S3). Even if we ignore this cancer-dependence and look at the structural types holistically, it is evident that the 3´ termini of 5´-tRFs and the 5´ termini of 3´-tRFs span a large number of choices (Figure 3). Notably, these preferences are very similar to what was reported recently for the plant *A. thaliana*^24^, which suggests common underlying biogenetic mechanisms and, possibly, functions. Furthermore, the cancer-dependence of the observed fragments suggests a tissue-specific dimension in the biogenesis of tRFs. This notion is supported, at least in part, by recent results showing that the channeling of tRNAs into the miRNA Dicer-Ago pathway depends on the structure of the RNA molecule^62^. Given that RNA folding is a dynamic process^63^, we posit that the observed differences in tRF cleavage patterns among tissues are caused by differences in each tissue’s molecular physiology.

The i-tRFs, a novel structural type that we discovered recently^4^, exhibit the largest diversity in TCGA. i-tRFs represent more than 75% of the 20,722 identified tRFs. As Figure 3 shows, i-tRFs have a multitude of preferred starting and ending positions. The choice of these endpoints strongly depends on cancer type (Supp. Table S3).

Despite the pronounced dependence of tRF profiles on cancer type, some isoacceptors stand out by producing tRFs with profiles that remain exceptionally consistent in healthy and diseased tissues, and across all cancer types. Of note here is the nuclear tRNA^HisGTG^ that produces -1U 5´-tRFs with lengths that range between 16 and 22 nt and have abundances that are characterized by a unique property. Specifically, the abundances of -1U 5´-tRFs with 3´ termini that differ by a single nt (all these tRFs share the same 5´ terminus) alternate between high and low, whereas their ratios remain constant across all analyzed normal and cancer samples, and all 32 cancer types. The resulting ‘see-saw’ pattern spans a limited and persistent range of ratios that can be seen in Figure 4 and Supp. Figure S3. It should be stressed, however, that even though the *ratios* of these -1U tRFs remains constant, their *absolute abundances* do change from cancer type to cancer type. We did not find any other isoacceptors whose tRFs exhibited this unusual behavior. The exquisite stability of these ratios across tissues, and the uniqueness of tRNA^HisGTG^ in this regard among tRNAs, leads us to conjecture that these -1U 5´-tRFs participate in fundamental cellular processes that are currently unknown.

Of equal importance is the finding that across all human tissues that we examined, the 5´-tRFs from tRNA^HisGTG^ contain primarily a uracil at the His(-1) position. This is a new and unexpected finding, because the mature tRNA^HisGTG^ requires guanylation of its 5´-terminus before it can be recognized by its cognate aminoacyl tRNA synthetase. By comparison, the levels of 5´-tRFs from tRNA^HisGTG^ with G, A, or C at the -1 position were low. Recent work with the human breast cancer cell line BT-474 suggests that -1U 5´-tRFs from the tRNA^HisGTG^ locus arise from the mature tRNA^28^. However, it is not clear for the time being whether the -1U 5´-tRFs that we discovered in the multitude of human tissues that we analyzed above arise from the processing of the mature tRNA^HisGTG^ or its precursor. Further complicating this determination is the fact that the DNA template at four of the 12 genomic loci encoding isodecoders of tRNA^HisGTG^ contains a T at the -1 position.

Given the nascent nature of this field and the apparent diversity and context-specific nature of the tRFs, it is not surprising that very little is known currently about their functional roles. With that in mind, we placed particular emphasis on leveraging the TCGA datasets to shed as much light as possible on this question. First, we found pairs of tRFs that are correlated across samples. Within a given cancer type, the same tRFs were found correlated across all available samples. However, different groups of tRFs were correlated across cancers (Supp. Figure S5B). We then extended this analysis to protein-coding transcripts and found a very rich repertoire of negative correlations involving tRFs and specific mRNAs. These tRF-mRNA anti-correlations depended strongly on cancer type (Supp. Figure S6E). Earlier reports by others and us provided evidence of tRFs acting like miRNAs via Argonaute loading^4,10,29^. In light of this, we also identified the group of mRNAs that are negatively correlated with miRNAs. The miRNA-mRNA anticorrelations depended strongly on cancer type as well (Supp. Figure S6E).

We wish to stress one point here. It is entirely possible that direct molecular coupling drives some of the uncovered correlations. However, in the absence of any additional information, it will be prudent to treat these relationships as associations. For example, these associations could result from a common upstream regulator, from belonging to the same pathway, or because some tRFs arise from the same precursor transcript. Considering the apparent diversity across cancer types, it appears that it will be necessary to unravel the mechanisms underlying the correlation patterns separately for each cancer. Moreover, the presented analysis makes it evident that tRFs have tissue-specific roles that are also more diverse than those of miRNAs (see below). Regarding the diversity in function, Ago-loaded tRFs are but one of multiple facets of tRF biology. Indeed, one should also recognize the interaction of tRFs with other RNA binding proteins and with the translation machinery^1,5,64^.

Even though the *specific mRNAs* that are found associated with tRFs *differ* between cancer types, the *pathways* to which these mRNAs belong show striking similarities across cancers. This observation is supported through a DAVID analysis of gene ontology terms that reveal four super-groups: cell adhesion, chromatin organization, and developmental processes (Supp. Figure S7D). Additionally, our analysis generated several observations that were reported recently in the literature. For example, we found several correlations involving tRFs from isoacceptors of Gly, Asp, Glu, and Tyr with the mRNAs of HMGA1, CD151, CD97 and TIMP3: these mRNAs were recently reported to be controlled by tRFs from these tRNAs in a YBX1-dependent manner^64^. Additionally, we enumerated more than 3,000 correlations of tRFs with ribosomal proteins, either mitochondrial or cytoplasmic, as well as more than 100 correlations of tRFs with aminoacyl tRNA synthetases, particularly IARS and MARS, which is in agreement with previous work in the field^5,65^.

We examined at the isoacceptor level the correlations of the above-mentioned four super-groups of mRNAs with tRFs. We split the mRNAs into those that are negatively correlated with tRFs only and those that are negatively correlated with both tRFs and miRNAs in the same cancer type. In each case, we computed the fraction of the 32 cancer types supporting a specific “tRF isoacceptor GO term” or a specific “miRNA+tRF isoacceptor GO term” relationship. This revealed a tight coupling of specific isoacceptors with specific GO categories that persists across multiple cancer types (Figure 5A), but is manifested by different tRF-mRNA pairings in each cancer (Supp. Figure S6E). A notable result of this analysis was that three of the 22 *mitochondrial* tRNAs (tRNA^LeuTAA^, tRNA^ValTAC^, tRNA^ProTGG^) were found to be negatively correlated with *nuclear* mRNAs in all four super-groups and in virtually all 32 cancers. This suggests the existence of a previously unrecognized tight coupling between MT and nuclear processes.

The seeming diversity of negatively-correlated tRFs and mRNAs in the face of persistent relationships between tRFs and pathways made us examine the cellular localization of proteins whose mRNAs are negatively correlated exclusively with either miRNAs or tRFs. When we looked across all cancers, we found a striking dichotomy (Figure 5B). Specifically, the proteins whose mRNAs are negatively correlated with miRNAs localized equally frequently in all of the considered destinations, and in virtually all cancers. On the other hand, the proteins whose mRNAs were negatively correlated with tRFs showed a preference for localization to the nucleus, cytoplasm, or cell membrane as well as a strong dependence on cancer type.

It is important to note here that, by comparison to miRNAs, the mechanisms of biogenesis and function of tRFs remain poorly understood for the most part. Nonetheless, as we mentioned above, it is known that short tRFs are loaded on Argonaute and act like miRNAs. With that in mind, let us assume for the moment that the uncovered anti-correlations imply tRF-mediated regulatory events that mirror the action of miRNAs on mRNAs. Then, our findings (Figure 5A) suggest an intriguing “division of labor” where some mRNAs are associated, and presumably regulated, solely by miRNAs, some solely by tRFs, and some by both miRNAs and tRFs. This synergistic hypothesis is further supported by the findings that are summarized by Figure 5B and indicate that tRFs are likely involved in cell-type-dependent interactions, analogously to what we reported previously for miRNAs^66^. An instance of this dynamic and contextdependent network of interactions was shown for tRFs from tRNA^Gln^ that interact with YBX1 in breast cancer cell lines^64^ but not in cervical cancer cell lines^65^.

Earlier^30-32^ and more recent work^15^ on the non-random placement of repeat elements on the genome as well as the finding that repeat elements become demethylated as stem cell differentiation progresses^67^, led us to examine one more possibility. Specifically, we examined possible associations between tRFs and the sequence composition of the genomic loci for mRNAs participating in these identified positive and negative correlations. Our analysis revealed intriguing associations between mRNAs that were negatively correlated with tRFs, and the “repeat-element content” of their respective genomic regions. In particular, we found that in many cancers, the genomic span of mRNAs that are positively correlated with tRFs are depleted in many categories of repeat elements. Analogously, in many cancers, the genomic spans of mRNAs that are negatively correlated with tRFs are enriched in many categories of repeat elements. In both cases, the observation holds true for instances of these repeats that are sense as well as antisense to these regions. The fact that the observed enrichment/depletion is highly statistically significant (p-val ≤ 0.001) and holds true for most of the 32 cancers suggests that these wide-ranging sequence relationships are likely being leveraged by a cancer cell’s post-transcriptional regulation layer^68^.

We note that several of the GO terms that are part of the four general pathways we described above (cell adhesion, chromatin organization, and developmental processes) are significantly over-represented in the group of genes that overlap with Alu elements^32^. Our results on the link of tRFs with repeat elements come on the heels of two recent and related publications. First, tRFs from the tRNA^GlyGCC^ isoacceptor were shown to repress expression of genes associated with the retroelement MERVL in mice^15^. Second, tRFs were shown to increase in *Arabidopsis* pollen in a Dicer-dependent manner and to specifically target transposable elements^69^. It is unclear currently whether the tRFs in human cancers act in a way similar to what is suggested in plants, i.e. to suppress transposon activity. Notably, the fact that tRFs have different correlations with repeat elements in different cancer types suggests a complex systemwide interaction network and a compendium of associated molecular events that differ from cancer type to cancer type. These correlations and data could start shedding light on the peculiar roles of repeat elements in human diseases and cancers^70^.

Previously, we demonstrated for miRNA isoforms that their abundance profiles in human tissues depend on a person’s sex, population origin, and race^71^, as well as on tissue, tissue state and disease subtype^72^. We also demonstrated that miRNAs are not unique in this regard and that tRNA fragments have the exact same dependency on sex, population origin, race, tissue, tissue state and disease subtype^4,9^. Working with the TCGA samples we had the opportunity to evaluate the possibility that similar dependencies might exist across all samples of a disease (independently of subtype) or across all samples of a fixed subtype. In this regard, we provided two characteristic examples. In the case of LUAD, we highlighted a dependency of tRF profiles on sex (Figure 6). In the case of the triple negative subtype of BRCA, we highlighted a dependency of tRF profiles on the patient’s race (Supp. Figure S8). Considering the emergence of tRFs as regulatory molecules in their own right, such dependencies are expected to modulate the regulatory events underlying a given disease in ways that have not been previously considered.

Considering the multitude and diversity of the uncovered tRFs, and the multiplicity of associations between tRNA fragments and various cancers, it is reasonable to assume that a lot more work will be required before the community can improve its understanding of the roles of tRFs in the cancer context. To facilitate investigations, we enhanced our MINTbase repository^22^ with a module that is specific to TCGA. The module provides access to all of the tRFs that we mined from TCGA. Importantly, the module permits very involved interactions with the contents of MINTbase by allowing elaborate search requests that require only minimal effort on the part of the user. We stress that although the TCGA portion of MINTbase is *static*, its non-TCGA portion is *dynamic* and growing steadily through the contributions of tRF profiles by different research teams. We designed the TCGA module in a way that permits users to compare TCGA findings with the ever-growing non-TCGA data.

In summary, analysis of the entirety of the TCGA repository revealed a very rich population of tRNA fragments. The identities and relative abundances of these fragments depend on cancer type. They also depend on the identity of the parental isoacceptor. Yet, tRF profiles remain essentially constant within samples of the same cancer type, underscoring the constitutive nature of these fragments. These tRFs exhibit strong associations with one another and with other molecular types such as mRNAs (and, by extension, miRNAs) suggesting the existence of numerous regulatory interactions that await discovery and characterization.

## METHODS

### Datasets

11,198 short RNA datasets were downloaded on October 16, 2015 from TCGA’s Cancer Genomic Hub (CGHub). We used datasets from both normal and tumor samples, which are identified by their TCGA barcode tag (01A, 01B and 01C for tumor; 11A, 11B and 11C for normal). These datasets already had adaptors trimmed and were converted back to FASTQ format using BamUtil’s *bam2FastQ* tool (http://genome.sph.umich.edu/wiki/BamUtil version 1.0.10). For each of these datasets, tRF profiles were generated using MINTmap^60^ and default settings. These profiles have been incorporated in MINTbase^22^.

Our analyses focused exclusively on whitelisted datasets. Generally, non-whitelisted samples are marked for withdrawal by the various TCGA projects for reasons that range from incorrect pathologic diagnosis to exclusion on the basis of patient medication history. Clinical metadata were downloaded from TCGA’s data portal on October 28, 2015. To help eliminate problematic and outlier samples that were identified by the various TCGA working groups, only datasets that did not have any special annotation notes within the clinical metadata were included (n=10,274).

### The various categories of tRFs

In terms of structural type, the tRFs overlapping a mature tRNA sequence fall in one of five possible categories^1,60^: a) 5´-tRNA halves or ‘5´-tRHs’^5,6,73,74^; b) 3´-tRNA halves or ‘3´-tRHs’^2,75,76^; c) 5´-tRFs^2,75,76^; d) 3´-tRFs^2,75,76^; and, e) the “internal tRFs” or “i-tRFs” that we discovered and reported recently^4^.

In terms of genomic origin, we characterize tRFs as “exclusive” or “ambiguous.” The sequences of exclusive tRFs are encountered only within the span of mature CCA-containing tRNAs, and appear nowhere else on the genome. Ambiguous tRFs on the other hand have sequences that can be found both in mature tRNAs (the “tRNA space”) and elsewhere on the genome. We recently published a methodology and standalone tool that automates the mining of tRFs from human RNA-seq datasets and automatically tags them as exclusive or ambiguous^60^. Our analyses were based on both exclusive and ambiguous tRFs.

In terms of length, we generated tRFs with lengths between 16 and 30 nt inclusive. It is important to note here that the short RNA-seq profiles for the samples of the TCGA repository were generated by running deep-sequencing PCR for 30 cycles. Although adequate for miRNAs, 30 PCR cycles will generate inaccurate profiles for those tRFs that are longer than 30 nt. In the various TCGA datasets, these longer tRFs appear truncated and, thus, are represented in the TCGA as “30-mers.” Our parallel work^7^ as well as our previous analyses of TCGA from BRCA subtypes^4^, and of non-TCGA datasets from prostate cancer^8^ and liver cancer^22^ show that there exist many distinct tRFs with length > 30 nt that are very abundant. Moreover, in the case of TCGA BRCA, we found that the “30-mer” tRFs are *differentially* abundant between normal breast and BRCA^4^, suggesting an association with disease states. We note that the adapter cutting step may have shortened artificially long tRFs into “30-mers”, a problem that does not arise when analyzing shorter molecules such as miRNAs^77^. Most of the analyses described below were based on tRFs with lengths 16-27 nt. Lest we miss potential important associations, we included tRFs with length 28-30 nt in those instances where doing so was warranted.

### Nomenclature and Notation

“Race” refers to a taxonomic rank below the species level, a collection of genetically differentiated human populations defined by phenotype. We adhere to the following NIH/TCGA designations: White (Wh) refers to person with origins in any of the original peoples of the far Europe, the Middle East, or North Africa; and Black or African American (B/Aa) refers to persons with origins in any of the black racial groups of Africa. Based on the provided information, the majority of TCGA samples are from either Wh or B/Aa donors. Smaller groups of samples were obtained from donors who are: a) American Indian or Alaska Native (i.e., persons having origins in any of the original peoples of North and South America, including Central America, and who maintain tribal affiliation or community attachment), b) Asian (i.e. persons having origins in any of the original peoples of the Far East, Southeast Asia, or the Indian subcontinent including, e.g., Cambodia, China, India, Japan, Korea, Malaysia, Pakistan, the Philippine Islands, Thailand, and Vietnam); and, c) Native Hawaiian or other Pacific Islander (i.e., persons having origins in any of the original peoples of Hawaii, Guam, Samoa, or other Pacific Islands). We named the tRFs using the scheme we introduced in our previous work^4^. Briefly it has two components separated by the @ sign, the first one being the mother tRNA and the second describing the coordinates within the mother tRNA (see Supp. Text for a more detailed description). As now we work with tRFs that are not exclusive to tRNA space, we append the string “Out” at the end of the name to indicate that the sequence of the tRF is also found outside of tRNA space.

### tRNA cleavage patterns

For each of the 32 cancer types, we examined the following attributes:

– location within the mature tRNA of the tRFs’ 5´ termini;
– nucleotide composition within a rolling dinucleotide window that surrounds the 5´ termini (positions 2/-1, -1/5´terminus, 5´-terminus/+1, +1/+2);
– location within the mature tRNA of the tRFs’ 3´ termini;
– nucleotide composition within a rolling-dinucleotide window surrounding the 3´ termini, as above; and,
– location of the tRFs’ 5´ and 3´ termini with respect to the mature tRNA endpoints and upstream-stem or downstream-stem of the nearest loop (D, anticodon, or T), as applicable.

Support for each of the attributes was calculated using tRFs above threshold. For each attribute, and for the cancer being studied, we calculated its normalized support in two ways: a) by considering only *distinct* tRF sequences ignoring their abundance; and, b) by repeating the analysis taking into account the abundance of the tRFs.

Supp. Figure S3 lists the complete set of histograms for all of the attributes that we tracked and all tRF categories. In addition to showing the results for each of the 32 cancers, we also provided histograms that combine the findings from all 32 cancer types.

### NMF analyses

The TCGA working groups have been making great use of non-negative matrix factorization^78^, or NMF, to cluster in an unsupervised manner the microRNAs (miRNAs) in the generated RNA-seq datasets. For this study, we replicated the NMF approach pioneered by the TCGA working groups leveraging tRF profiles (instead of miRNA profiles). We ran NMF (with R’s NMF module, version 0.20.6) in an unsupervised manner using the top 30% most variable tRFs that passed Threshold-seq and had mean RPM >=1. Only tRFs with lengths 16-27 nt inclusive were used in these analyses. For each cancer type, the input used during clustering was a matrix comprising the RPM-normalized tRF profiles of the whitelisted datasets (see above) for the cancer type. Only the tumor datasets of each cancer type were used. For SKCM, NMF clustering was carried out separately on the primary tumor and the metastatic samples. Silhouette widths were generated from the final NMF consensus membership matrix (n=500 iterations per run). NMF^79^ was run using values of *k* ranging from 2 through 10 inclusive. For GBM, NMF clustering was not carried out because of the small number (5) of available datasets.

### Correlation analyses

For each cancer type separately, we first filtered the tRFs and the genes based on their abundance. For this step, we considered all tRFs and all miRNAs with a median expression ≥ 1 RPM and the genes (TCGA’ s *rsem_genes.normalized_results* files) with an average expression of ≥ 1 RSEM. We applied an additional filter for mRNAs, keeping only the top 50% most expressed entities. Then, we computed all pairwise tRF-tRF Spearman correlation coefficients, as well as all tRF-mRNA and all miRNA-mRNA Spearman correlation coefficients for all expressed genes. We corrected the P-values to FDR scores, using the *p.adjust* function in the R base package with the method argument ‘FDR.’ We only kept correlation coefficients that had an FDR ≤ 0.01 and an absolute value larger than 0.333. For the sex- and race-specific networks, we relaxed the FDR threshold to 0.05. In those instances where several thousands of coeffi-cients survived these thresholds, we kept only the top (for positive correlations) or bottom (for negative correlations) 5,000 correlations. Computations were done using python and the *numpy* (version 1.11.1) and *scipy* (version 0.18.1) packages.

Probabilities for the tRF-tRF networks were computed as the number of nodes that satisfy the respective criteria divided by the total number of nodes in each network.

Pathway analysis was run separately for the collection of genes that were negatively correlated with a) tRFs, or b) miRNAs. Specifically, DAVID (version 6.8)^80^ was run with these two collections of genes and the overlap with GOTERM_BPFAT, GOTERM_MFFAT, GOTERM_CCFAT, and, KEGG_PATHWAY terms was calculated and filtered at an FDR threshold of 5%. The genes that were used in the correlation analysis in each cancer served as the background gene list for the DAVID tool.

### Protein localization

Information on protein localization was downloaded from UniProt^81^ and only the manually reviewed human proteome (queried on November 27, 2016) was used. For each cancer type and correlation group (positive or negative), the distribution of the localization of gene products was computed as a percentage in each of the following cellular compartments: Nucleus, Cell membrane, Mitochondrion, Endoplasmic Reticulum and Golgi apparatus (ER/Golgi), Cytosol, Organelles (peroxisomes, endosomes, lysosomes) and extracellular proteins (marked as “Secreted”). Gene products that are not part of any of these categories, have an unknown localization, or do not have a matching UniProt entry were assigned to the “Other” category: this category comprised, on average, 17% of the gene products across cancers. To estimate the statistical enrichment or depletion, we performed Monte-Carlo simulations with 10,000 iterations and built the expected distribution for the mRNA’s product localizations by randomly selecting the same number of mRNAs in each iteration. We performed the simulation separately for each cancer type and for each nuclear tRFs, mitochondrial tRFs and miRNAs. The results are presented as *enrichment* (yellow color) or *depletion* (purple color) for each compartment, calculated as a Z-score of lower than -2 or greater then +2 with respect to the expected distributions.

### Overlap with RepeatMasker entries

To calculate the overlap with RepeatMasker (http://www.repeatmasker.org;hg19version4.0.5) elements, the union of genomic regions of all splice variants of a gene was taken in order to capture repeat elements that are specifically localized downstream of the transcription start site^15,32^. Then, we counted how many of these genomic regions overlapped on the sense or antisense strand with each one of the repeat families of Repeat-Masker. In order to evaluate whether this ‘observed’ overlap corresponded to enrichment or depletion of repeat elements, we ran Monte-Carlo simulations to create an ‘expected’ distribution of overlap with RepeatMasker elements. In more detail, in each of 10,000 iterations we randomly selected from the total pool of genes included in the correlation analysis (all the expressed genes that passed the expression filtering) the same number of genes as the number of unique genes that were correlated with tRFs. For each cancer type, positive and negative correlations were analyzed separately (a total of 64 simulations each with 10,000 iterations). After each iteration, we calculated the overlap with RepeatMasker elements as described above and used it to create the ‘expected’ distribution. Based on this distribution (normal distribution), we calculated the z-score of the observed enrichment for each repeat family.

### Disambiguation of the genomic origin of tRFs

To investigate whether non-exclusive tRFs as well as nuclear tRFs are enriched or depleted in our correlation analyses, we performed Monte-Carlo simulations analogously to the way we calculated overlap with repeat elements. Specifically, we performed 10,000 iterations and in each one we calculated the ratio of non-exclusive tRF and of nuclear tRF based on a randomly chosen set of tRFs equal in size to the set of tRFs participating in the tRF-mRNA correlations. This was carried out separately for each cancer type. We then built a distribution of these ratios and calculated the enrichment or depletion for non-exclusive tRFs and for nuclear tRFs, independently per cancer type.

### Multivariate statistical analysis and data visualization

Hierarchical Clustering and Principal Component Analysis, as well as network visualizations were run and plotted in R, as we previously described^4,66,72^.

## ACKNOWLEDGMENTS

We thank all patients who consented to have their samples analyzed by▭TCGA Network. We are indebted to the National Institutes of Health for making the TCGA data publicly available. We thank Dr. Eric Londin, Dr. Megumi Shigematsu, Dr. Takuya Kawamura, as well as the other members of the Computational Medicine Center for discussions and input on this manuscript. The work was supported partially by a William M. Keck Foundation grant (IR), by NIH/NCI R21-CA195204 (IR), and by Institutional Funds.

## AUTHOR CONTRIBUTIONS

IR, AGT, PL conceived and designed the study with contributions from RM, YK, and VP. IR supervised the study. PL downloaded the data and generated the tRF profiles. AGT, PL, RM, and IR mined the tRF profiles and analyzed the results. PL, IR, AGT and VP performed the cleavage pattern analyses. RM and IR analyzed the His(-1U) tRFs. PL generated the NMF clusters that were subsequently analyzed by AGT, RM, and PL. AGT and IR performed the correlation analyses. VP implemented MINTbase v2. IR, AGT and RM wrote the manuscript with contributions from PL, YK, and VP. All authors have read and approved the final manuscript.

